# Flexible belief updating drives the childhood advantage in statistical learning

**DOI:** 10.64898/2026.06.30.735487

**Authors:** Orsolya Pesthy, Eszter Tóth-Fáber, Cintia Anna Nagy, Márton Németh, Karolina Janacsek, Dezso Nemeth

## Abstract

Children often outperform adults in probabilistic statistical learning tasks, yet the mechanisms underlying this developmental advantage remain poorly understood. Here, we used eye-tracking measures of belief updating to examine how children and adults acquire and update predictions in a probabilistic sequence-learning task. Using the standard (oculomotor) reaction time measure, children showed stronger statistical learning than adults, replicating previous behavioral findings while revealing a more detailed profile of developmental differences in statistical learning. Critically, children updated their predictions more frequently: they were less likely to repeat previous predictions and more likely to shift their expectations in response to new input. Adults, in contrast, showed greater persistence, tending to maintain prior predictions even when those predictions were inconsistent with the underlying statistical structure. Despite these pronounced differences in updating behavior, the processing and use of prediction errors were remarkably similar across age groups. These findings indicate that developmental differences in statistical learning do not primarily arise from how prediction errors are computed, but rather from how prior beliefs and incoming information are weighted during belief updating. Children’s enhanced learning may therefore reflect reduced reliance on stable priors and greater sensitivity to current sensory evidence, supporting a more exploratory learning strategy. Adults, by contrast, appear to favor an exploitative strategy that stabilizes existing predictions but reduces flexibility in probabilistic environments. More broadly, the results suggest that developmental changes in statistical learning may reflect age-related differences in how readily learners revise their predictions in response to incoming evidence. By integrating sensitive oculomotor measures with analyses that probe the mechanisms underlying belief updating, the present study provides a more fine-grained account of how predictive learning changes across development and offers a framework for reconciling previously inconsistent developmental findings in statistical learning.

## Introduction

The brain continuously extracts environmental regularities to build internal predictive models without relying on explicit instructions or rewards. This capacity, known as statistical learning, drives the acquisition of complex behaviours from infancy through old age, underpinning language development, motor habits, and social interactions (Fiser & Aslin, 2001; Kaufman et al., 2010; Saffran et al., 1996; Ullman, 2004). Yet, while this core learning mechanism is remarkably robust (Arciuli & Simpson, 2012; Kobor et al., 2017; Romano et al., 2010), its developmental trajectory remains heavily debated. To resolve these seemingly contradictory developmental trajectories, the field requires a mechanistic account of how these unsupervised, reward-free models evolve. Specifically, we must understand how learners compute prediction errors — the mismatch between their predictions and actual sensory input — and update their prior beliefs across development. Viewed through this computational lens, cognitive maturation is not a uniform change in capacity, but an adaptive shift in information processing strategy. Development marks a transition from a higher level of information seeking and broader, more exploratory processing style to a more stable, knowledge-driven exploitation of the adult mind (Frankenhuis & Gopnik, 2023; Gopnik, 2020; R. Wu et al., 2016). Understanding developmental changes in exploratory behaviour, belief updating, and sensitivity to environmental statistics is therefore essential for explaining not only age-related differences in learning, but also the (qualitatively or quantitatively differing) mechanisms that foster them.

Previous research has revealed diverse developmental trajectories in statistical learning, largely influenced by the modality of the input and the nature of the underlying regularities (Forest et al., 2023). Statistical learning tasks employing auditory stimuli seem to consistently show age invariance when the stimuli are linguistic (François et al., 2013; Raviv & Arnon, 2018; Shufaniya & Arnon, 2018), and improvement with age when the stimuli are non-linguistic (Arciuli & Simpson, 2011; Raviv & Arnon, 2018; Schlichting et al., 2017). The picture is more complex in the visual domain, where findings are mixed (Bertels et al., 2014; Janacsek et al., 2012; Johnston et al., 2026; Jost et al., 2015; Juhasz et al., 2019; Lukács & Kemény, 2015; Nemeth et al., 2013; Raviv & Arnon, 2018; Shufaniya & Arnon, 2018; Tóth-Fáber et al., 2026). In this domain, results appear to depend heavily on the type of regularity, with a relatively consistent advantage for children in tasks with probabilistic regularities (Janacsek et al., 2012; Johnston et al., 2026; Juhasz et al., 2019; Nemeth et al., 2013; Tóth-Fáber et al., 2026), and either an advantage for (young) adults (Lukács & Kemény, 2015) or age invariance (Lum et al., 2010) in tasks with deterministic regularities. Collectively, these findings suggest that task demands strongly influence the developmental trajectory of statistical learning. Here, we applied eye-tracking and a novel analytical approach to disentangle the mechanisms underlying statistical learning. Understanding how internal models of environmental regularities are formed and updated may help explain why different tasks produce markedly different developmental patterns.

Uncovering the underlying mechanisms may provide the key to understand developmental changes in statistical learning, as children may rely on slightly different strategies when learning environmental regularities. One candidate mechanism is how prediction errors are processed: how much children and adults trust the discrepancies between their predicted and the actual outcome (i.e., the precision or weight of the prediction error). Most developmental evidence comes from reinforcement learning paradigms, showing mixed results: Some studies report lower error precision in children (Bruckner et al., 2025), or that adults exhibit stronger neural signs of prediction errors (Rodriguez Buritica et al., 2024), while others found no age differences (Javadi et al., 2014). Importantly, prediction errors appear to be computed similarly to adults even as early as in infanthood (Zhang et al., 2019). Yet, immature connectivity may alter how these signals influence behaviour across development (Van Den Bos et al., 2012). However, most of these studies used reinforcement learning tasks, thus, it remains unclear how prediction error processing develops in tasks without reward or explicit feedback such as statistical learning tasks.

Another related candidate mechanism that may show developmental changes is the dynamics of updating one’s belief. Evidence primarily from the reinforcement learning literature suggests that children tend to change their previous predictions more than adults, reflecting a more exploratory learning style (Jepma et al., 2020). This tendency may be linked to their more distributed attention (Blanco & Sloutsky, 2020; Frank et al., 2021) and broader information sampling (Wan & Sloutsky, 2025). Adults are better able to balance stable strategies with the exploration of new options (C. M. Wu et al., 2018), nevertheless, they tend to stick to their previous predictions more often than children (Constantino & Daw, 2015), at least in reward contexts where this exploitation is directed toward the most rewarding option (Blanco & Sloutsky, 2020, 2021, 2024). Here, we aimed to test whether differential tendencies to update or repeat previous predictions could give rise to developmental differences in the absence of reward, such as in statistical learning.

To this end, here, we used the gaze-contingent, eye-tracking version of the Alternating Serial Reaction Time (ASRT) task to introduce new statistical learning metrics. Using these metrics enables us to capture iterative updating, as well as the direct assessment of error processing in a reward-free probabilistic task. Eye-tracking provides fine-grained examination of underlying cognitive processes, including a closer approximation of prediction errors (Tal & Vakil, 2020; Vakil et al., 2021). We measured anticipatory saccades, that is, saccades in the response-to-stimulus interval that reflect expectations about the statistical structure of the task. This way, prediction formation and its violation can be tracked in real time (Hann et al., 2026). This approach offers advantages over visuomotor statistical learning tasks in which participants typically press buttons corresponding to stimuli, and errors are defined as mismatches between button presses and presented stimuli. Such operationalization of errors reduces the sensitivity in probabilistic contexts for two reasons. First, while in deterministic tasks, erroneous responses necessarily mean a deviation from the underlying regularity, in probabilistic environments, “erroneous” responses still can reflect an accurate representation of the task probability structure. For example, it is possible to select the highest-probability option but a low-probability event occurs (due to the probabilistic nature of the task). Second, correct button presses can mask cognitive-level prediction errors. Eye-movements are more automatic and more directly tied to prediction-related processing than overt responses, thus reflecting internal belief states more directly (Tal & Vakil, 2020). Here, we developed metrics that open a window to the participants’ error processing.

Furthermore, eye-tracking enables the examination of how participants update largely implicit prior beliefs about environmental regularities (Friston, 2010; Hann et al., 2026). In Bayesian terms, predictions are formed iteratively using prior beliefs (priors) and incoming sensory input (Friston, 2010). Because anticipatory saccades occur during the response-to-stimulus interval, with no relevant sensory input present, they provide a closer proxy of priors compared to reaction times or button-press responses (Hann et al., 2026). By tracking predictions on a trial-by-trial basis, we can directly observe how participants update their beliefs: specifically, whether and how their predictions change following a given event. Crucially, using this approach, we can quantify the impact of prediction errors on the updating process. Tracking the effect of errors on updating is particularly important in probabilistic tasks, where not all errors should contribute equally to belief updating. As noted above, some errors may reflect accurate knowledge of the underlying probability structure of the task. We therefore distinguish learning-dependent and not-learning-dependent errors. Learning-dependent errors occur when a participant predicts the highest-probability event but, by chance, a low-probability event occurs. In an optimal learner, such errors should not drive belief updating, as doing so would degrade an already accurate representation. In contrast, not-learning-dependent errors arise when a participant predicts a low-probability event; these errors should prompt updating, as they indicate a genuine mismatch between the participant’s beliefs and the task’s statistical structure.

This study compared adults and 8-13-year-old children on a gaze-contingent eye-tracking task involving probabilistic regularities, to clarify whether the two groups rely on similar or distinct mechanisms during statistical learning. Beyond the usual statistical learning measures such as (oculomotor) reaction times, we examined age differences captured by anticipatory saccades. Specifically, we used novel eye-tracking measures to compare how the two groups respond to errors measured by saccadic behaviour – which we consider a behavioural proxy of prediction errors (c.f. Hann et al., 2026). The probabilistic nature of our task enabled us to differentiate the effect of errors arising from insufficient knowledge of the statistical structure (not-learning-dependent error) versus those reflecting accurate knowledge in a probabilistic context (learning-dependent error). We also tested how participants update their prior beliefs following prior responses. By combining these metrics, this study provides a more comprehensive account of potential developmental differences in statistical learning, offering insights into whether children rely on qualitatively different mechanisms than adults and how such mechanisms may shape learning outcomes across development.

## Methods

### Participants

We recruited a total of 92 participants: 43 children between 8-13 years and 49 adults. We excluded three children because they reached 14 years by the day of the experiment. We further excluded four children and three adults because they did not finish the entire task, either due to calibration issues or fatigue, resulting in an 82-participant sample. We further excluded participants based on low eye-tracking data quality. Data quality was calculated by task epochs, that are analysis units of about 5 minutes of practice time (see the Procedure section for details). We excluded participants with a missing data ratio above 20% (two adults), a median distance from the screen outside the functional range of our device (50-90 cm; no participants), or outlier precision metrics (eye-to-eye root mean square: five children, three adults, sample-to-sample root mean square: three children, three adults) in any epochs. Please note that some participants may overlap in these data quality measures. This resulted in a final sample of 68 participants: 28 children (*M_age_* = 10.71 years, *SD_ag_*_e_ = 1.58 years, 15 girls, 13 boys), and 40 adults (*M_age_* = 23.05 years, *SD_ag_*_e_ = 3.07 years, 25 women, 15 men). All participants gave their informed written consent (in the case of children, written consent was given by their caregiver). Participants did not receive financial compensation. The study was approved by the Eötvös Loránd University’s Research Ethics Committee in Budapest, Hungary (2022/134) and was conducted in accordance with the Declaration of Helsinki. The general saccade metrics of the groups are shown in Table 1.

**Table 1.** Saccade metrics in children and adults.

| Group | Duration |  | Amplitude |  | Velocity |  |
| --- | --- | --- | --- | --- | --- | --- |
|  | Mean | SD | Mean | SD | Mean | SD |
| Adults | 33.58 | 18.848 | 2.841 | 3.09 | 84.84 | 55.74 |
| Children | 33.33 | 19.283 | 2.815 | 3.25 | 80.25 | 54.23 |
| Difference | $t(133760) = -2.36,$<br>$p = 0.018$ | | $t(133760) = -1.38,$<br>$p = 0.168$ | | $t(133760) = -14.92,$<br>$p < 0.001$ | |

### Task

To measure statistical learning, we used the eye-tracking version of the Alternating Serial Reaction Time (ASRT) task (Howard & Howard, 1997; Zolnai et al., 2022), see Figure 1. In this task, participants learnt a probabilistic sequence. Participants saw four empty circles (arranged in a square shape), and one of the circles turned blue. They were instructed to fixate on the blue circle as fast as possible. Participants took as much time as they needed to fixate on the blue stimulus (i.e., the task was self-paced). After the fixation, the given circle turned empty, and the next stimulus appeared with a 500 ms response-stimulus interval (RSI).

**Figure 1.**
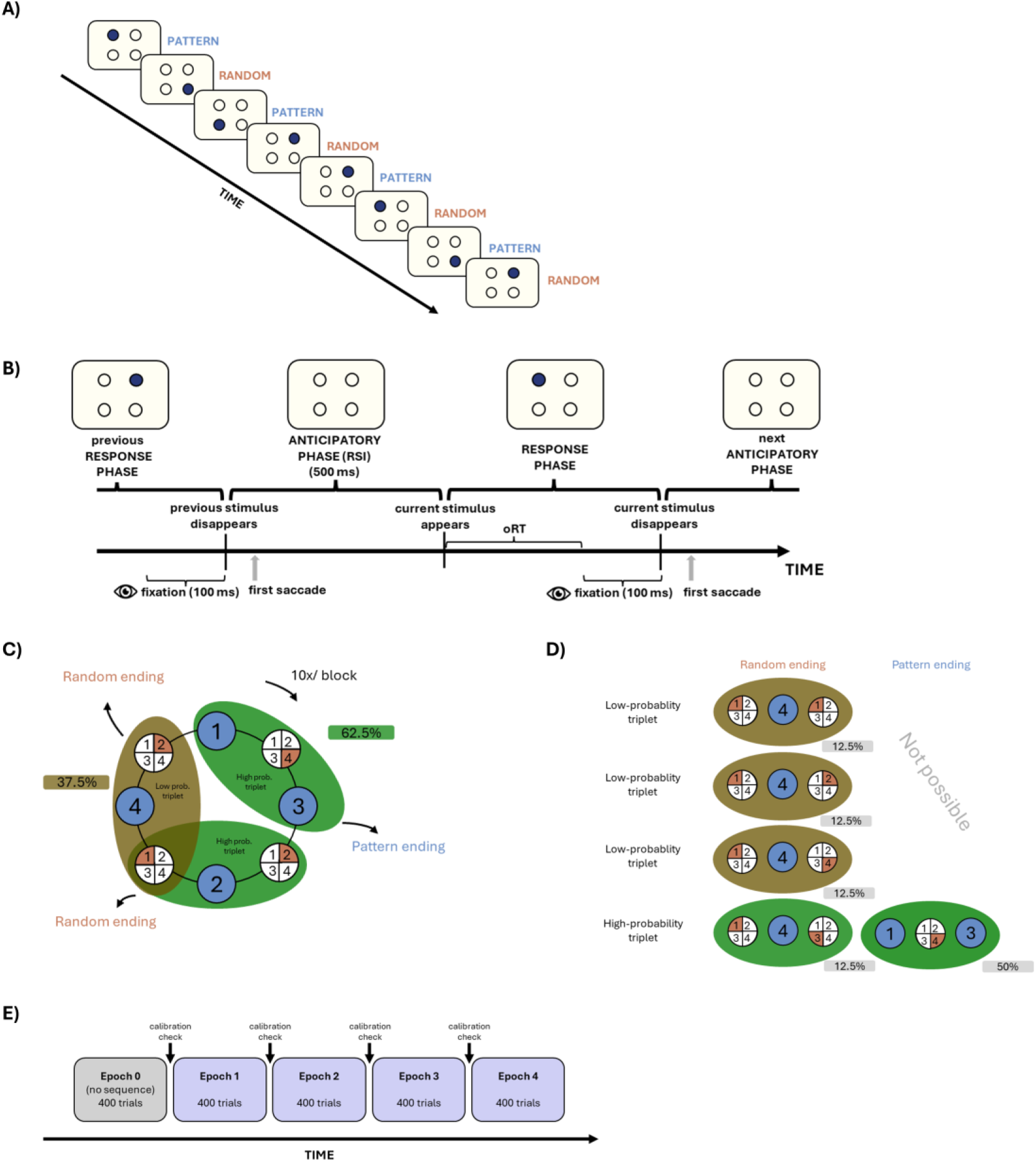
Structure of the oculomotor ASRT task and the experimental design. **A)** During the task, participants saw four empty circles arranged in a square on the screen, one of which turned blue during each trial. In the stimulus series, every second trial was part of an 8-element probabilistic sequence. Random elements were inserted between the pattern elements (e.g., 1-r-3-r-2-r-4-r, where the numbers indicate the position of one of the four circles on the screen and “r” indicates random positions). **B)** Each trial consisted of two phases: an anticipation phase (when the stimulus had not yet appeared on the screen) and a response phase. During each trial, one of the four empty circles turned blue. The participants’ task was to direct their gaze to the blue circle as quickly and accurately as possible and to maintain their gaze there for at least 100 ms. Once the fixation was adequate, the stimulus disappeared and the next stimulus appeared after 500 ms. The oculomotor reaction time (oRT) was defined as the time elapsed from the appearance of the stimulus to fixation on the stimulus location. **C)** Formation of triplets in the task. Pattern elements are marked in orange (they always appear in the same place during the task), while random elements are marked in blue (they appear randomly in one of the four possible locations). Each trial was categorized based on the third element of three consecutive trials (triplet). Due to the probabilistic structure, some triplets occurred more frequently (marked with a green background) than others (marked with a brown background). Statistical learning was defined as the decrease of reaction time for high-probability trials compared to low-probability trials. **D)** High-probability triplets can consist of two sequential and one random element, which occurs in 50% of trials; or two random and one sequential element, which occurs in 12.5% of trials. Overall, 62.5% of all trials constitute the last element of high-probability triplets, while the remaining 37.5% constitute the last element of low-probability triplets. **E)** The task was divided into blocks, each consisting of 80 trials (ten repetitions of a series of eight elements), preceded by five initial random warm-up trials, which were excluded from all analyses. A set of five blocks is called an epoch, which thus contains 400 trials. Participants were familiarized with the task by completing a practice epoch, which contained only random trials. The actual task consisted of four epochs containing the probabilistic sequence. After each epoch, calibration was checked based on the last 20 trials, and the eye tracker was recalibrated if necessary.

Unbeknownst to the participants, there was a hidden sequence in the task. Every second stimulus appeared in line with the sequence (pattern trials “P”) with random trials in between when any of the four circles could turn blue (random trials “R”) creating an 8-element long sequence e.g., 1-R-3-R-2-R-4-R, where the numbers represent one of the four circles on the screen, and “R” indicates a randomly selected one). Due to this alternating structure, certain triplets (three consecutive elements) were more probable to occur than others, so two types of triplets can be distinguished based on the probability of the last element of the triplet. High-probability triplets could be formed in two ways: either with two pattern trials and one random (P-R-P) or one pattern trial and two random (R-P-R) (e.g., 1-X-3, 3-X-2, 2-X-4, and 4-X-1 triplets, where “X” indicates either a pattern or random trial). In contrast, low-probability triplets could be formed only in one way: one pattern trial and two random (R-P-R) (e.g., 1-X-2 or 1-X-4, where “X” indicates a pattern trial). There was a total of 16 possible formations for high-probability triplets and 48 for low-probability triplets. Each high-probability triplet had an individual appearance probability of approximately 3.91%, while each low-probability triplet had an appearance probability of approximately 0.78%. This resulted in a total probability of 62.5% (16 × 3.91) for high-probability triplets and 37.5% (48 × 0.78) for low-probability triplets.

### Procedure

The task consisted of blocks, each containing 80 trials (ten repetitions of the 8-element sequence) and 5 warm-up random trials at the beginning (which was excluded from the analysis). After each block, participants saw their mean reaction time of the preceding block which they could use to improve their performance and had the opportunity to have a short self-paced break. Participants completed 5 practice blocks (which only contained random trials and served as a familiarization with the task and was not analysed) and then 20 blocks of the task. We used Tobii Pro X3-120 eye-tracking device (Tobii AB, 2017) to measure eye-movements and fixations. More details on the gaze position and fixation estimation can be found in (Zolnai et al., 2022). In the beginning of the task there was a calibration via Tobii Pro Eye Tracker Manager (Tobii Pro A. B., 2020), in which participants looked at dots, appearing in one of the four corners of the screen or in the middle, until it disappeared. Then we validated the calibration of the eye-tracker with 20 random trials. If there was no extreme reaction time, i.e., above 1000 ms, we deemed the calibration correct. If there were extreme reaction times, we repeated this procedure until reached zero extreme reaction time. If we failed through six calibrations, we stopped the experiment and excluded the participant. After every fifth block, we checked the calibration of the eye-tracker based on the last 20 trials of the block and recalibrated if necessary.

### Eye-tracking data quality control

To ensure eye-tracking data quality, we excluded participants with at least one outlier epoch in any of the quality measurements. The calculated measurements were the following: missing data ratio, eye distance from the screen, and precision.

The percentage of invalid samples for each epoch was calculated for each participant, excluding those with 20% or higher missing data ratio (Hessels et al., 2020). Data loss can be due to various reasons such as blinking, eyelashes, eye tracker failure, etc. The participant’s median distance from the screen epoch was also calculated: participants were excluded if their distance was outside of the device’s functional range (50-90 cm). Precision is defined as the consistency of the measured gaze direction when the participant is looking at the current stimulus. We calculated two types of scores to evaluate precision. First, we calculated the root mean square of sample-to-sample which shows the root mean square differences between consecutive samples during a fixation, that is when we can assume that the participant is looking at the same point. We calculated the median sample-to-sample root mean square for each participant’s epoch. We also computed the root mean square of the eye-to-eye distance. This involved calculating the positional differences between the two eyes for each sample within a fixation window (excluding samples lacking valid data for both eyes) and deriving the root mean square value of these differences. Outlier epochs were defined using boxplots, that is greater than the upper bound of the 1.5 interquartile range or less than the lower bound of the range. The expected and observed data for each measurement and the number of excluded epochs can be seen in Table 2.

**Table 2.**
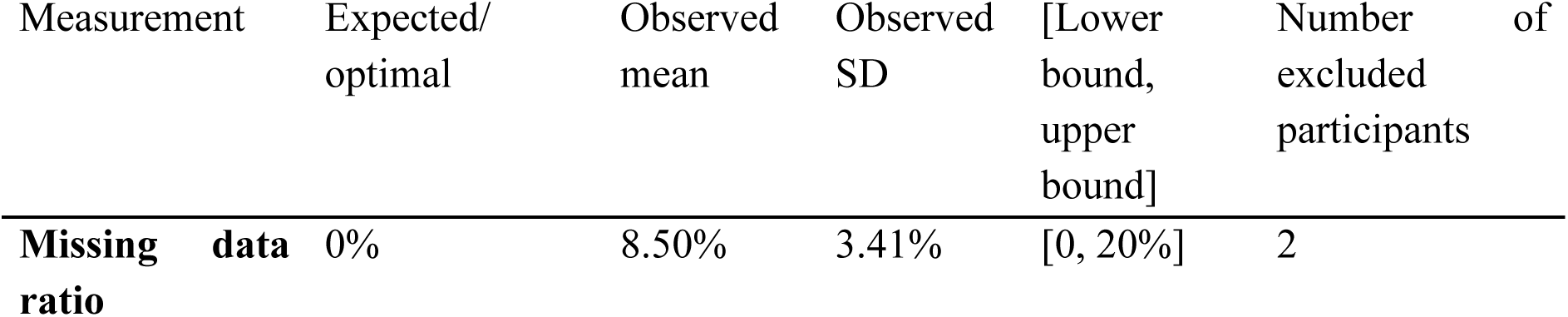
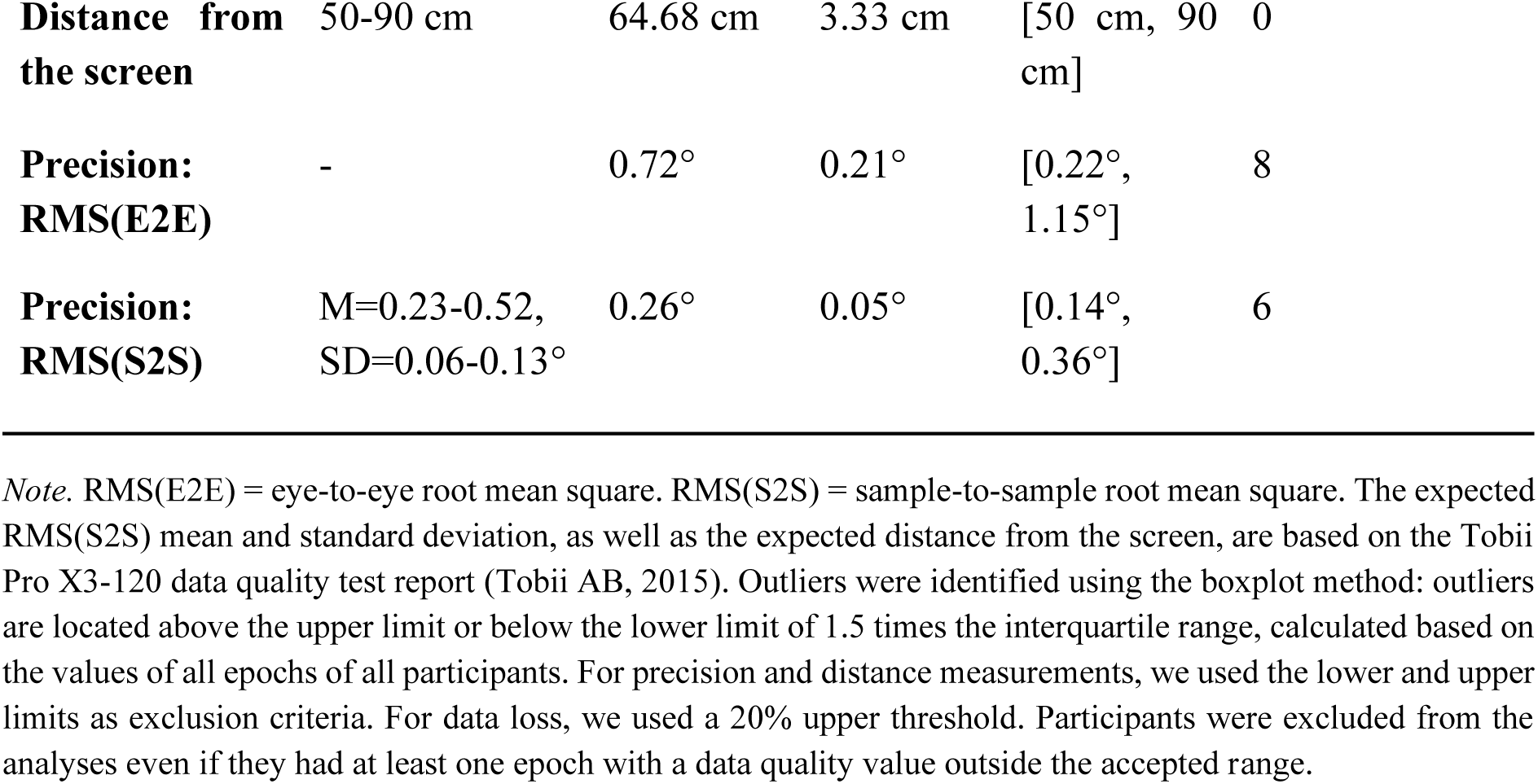
Data quality measures of eye tracking data.

### Data preprocessing and saccade identification

For preprocessing the eye position coordinate time series, we used Python (version 3.13), applying the *numpy* (version 2.4.1), *pandas* (version 3.0.0), and *SciPy* (version 1.13.1) packages (Harris et al., 2020; Reback et al., 2020; Virtanen et al., 2020). For all data preparation and analysis codes, see https://osf.io/qm8av/.

First, we calculated the average eye position coordinate of the two eyes using a hybrid method in order to minimize missing data: where both eyes’ data were available, we calculated the mean of their x and y coordinates. We used a single eye position coordinates when the other eye position coordinates were unavailable. Where both eyes’ coordinates were missing, the sample was marked as missing data.

We preprocessed the data for oculomotor reaction time (oRT) analyses using the algorithm described in Zolnai et al. (2022). We defined oRTs as the time interval between stimulus onset and the start of the first fixation on the location of the given stimulus. Fixations were defined as at least 100 ms periods where the gaze remained within a 4×4 cm square around the stimulus location.

We identified saccades using the procedure described in Hann et al. (2026). Potential saccade onsets and offsets were identified by computing the sample-to-sample eye velocity (in degrees of visual angle/s). We applied a Savitzky-Golay filter on the velocity time series (frame length: 7 samples, polynomial order: 5) (Holmqvist et al., 2012; Lum, 2020) to reduce high-frequency noise stemming from eye-tracker imprecision, via the *savgol_filter* in the *scipy.signal* package. A sample was classified as a potential part of a saccade if its velocity exceeded 30 °/s, but remained below 1000 °/s to ensure biological feasibility. Then, only candidate saccades that showed a saccade-like profile were retained: saccades had to first reach the 30 °/s threshold, then peak at minimum 50 °/s, followed by dropping under 30 °/s again. After this filter, the potential saccades underwent an examination of duration (in ms) and amplitude (in degrees of visual angle). To erase artifacts, candidate saccades with durations over 100 ms or amplitudes exceeding 30° were removed. Due to the extraction method described above, the minimum detectable saccade duration was three samples (onset, peak, offset), corresponding to 25 ms, as our device had a 120 Hz sampling rate.

Following the saccade extraction, we computed the estimated saccade target, and classified it into one of the possible stimulus locations to estimate potential predictive saccades. We computed saccade vectors in visual-angle space between the gaze position at the onset and offset of each saccade (saccade vector). This vector was then compared to the vectors between the saccade onset and each stimulus location (stimulus vectors). The stimulus vector which formed the smallest angle with the saccade vector was labelled as the predicted stimulus, and the angle of the saccade vector and predicted stimulus vector was saved as the predictive angle. We excluded saccades with a predictive angle of 90° or greater from all further analyses, as these indicate eye movements that leave the task-relevant area of the screen (i.e., they move toward the edges of the screen). See Figure 2 for visualization.

**Figure 2.**
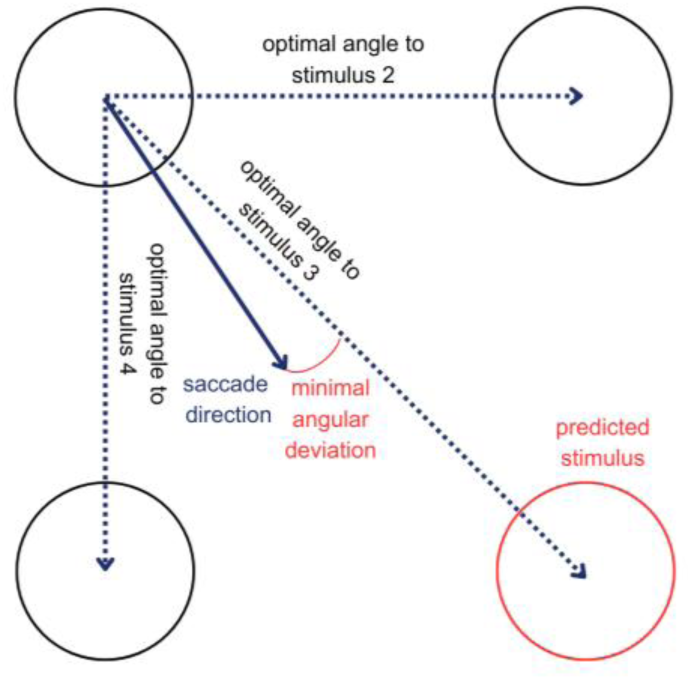
Determining predicted stimulus. A saccade vector (solid blue arrow) was defined by the gaze coordinates at the onset and offset of the saccade. From the onset position, reference vectors (dashed black arrows) were drawn to the centres of all possible stimulus locations. We then calculated the angular difference between the saccade vector and each reference vector. The stimulus location associated with the smallest angular deviation (red arc) was taken as the predicted target stimulus.

Lastly, we retained only the first saccade of each trial’s response-to-stimulus interval, as they are reported to reflect predictive processes (Jiang et al., 2014). Trials in which no such saccade was detected were marked as missing.

### Statistical analysis

Data preprocessing was performed in Python 3.13, using NumPy (version 2.4.1, Harris et al., 2020), pandas (version 3.0.0, Reback et al., 2020), SciPy (version 1.13.1, Virtanen et al., 2020). Linear and generalized linear mixed models were fitted with R (version 4.5.1), using the *mixed* function of the *afex* package (Singmann et al., 2024). We used the *emmeans* package (Lenth, 2024) for post hoc contrasts. Degrees of freedom were obtained using the Satterthwaite approximation (Satterthwaite, 1941). Figures were made in Python, using *matplotlib* (version 3.9.2) and *seaborn* (version 0.13.2) (Hunter, 2007; Waskom et al., 2017). We used an alpha level of 0.05 for all analyses. For the oRT analysis, we excluded trials that belonged to a repetition (trials that appeared in the same location as the previous trial) as participants show pre-existing tendency to react differently on these trials (Soetens et al., 2004). For testing hypotheses, we ran linear-mixed models. The final model was selected through the following procedure. First, we specified the full random structure, including both random slopes and intercepts. If this model failed to converge, we removed the interaction between random effects. If convergence issues persisted, we sequentially excluded the random effect that explained the least variation, continuing this process until the model reached convergence or until we arrived at the minimal random structure, which included only random intercepts for each participant (Barr et al., 2013; Bates et al., 2015). The *p* values of all post hoc tests were corrected using the Holm method for multiple comparisons (Holm, 1979). All *p* values were two-tailed. For a summary of all metrics, see Table 3.

**Table 3.**
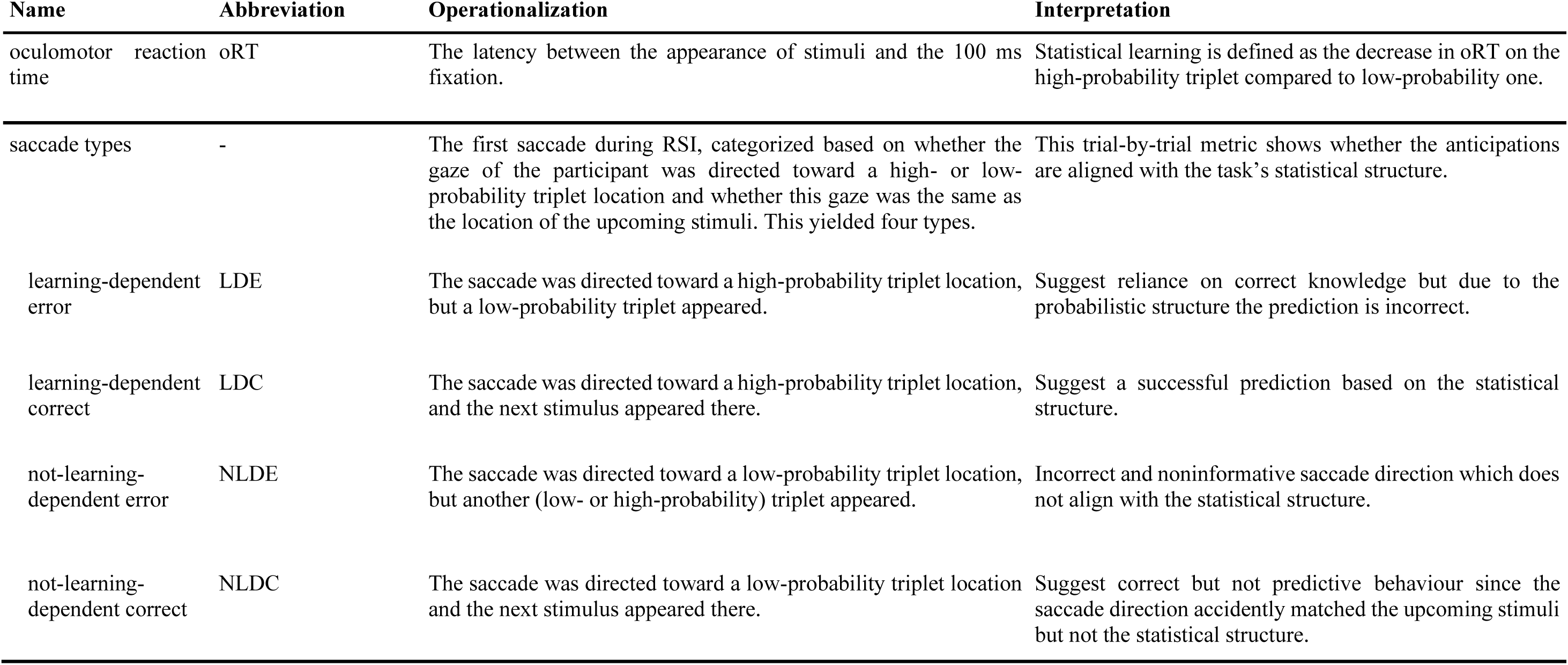
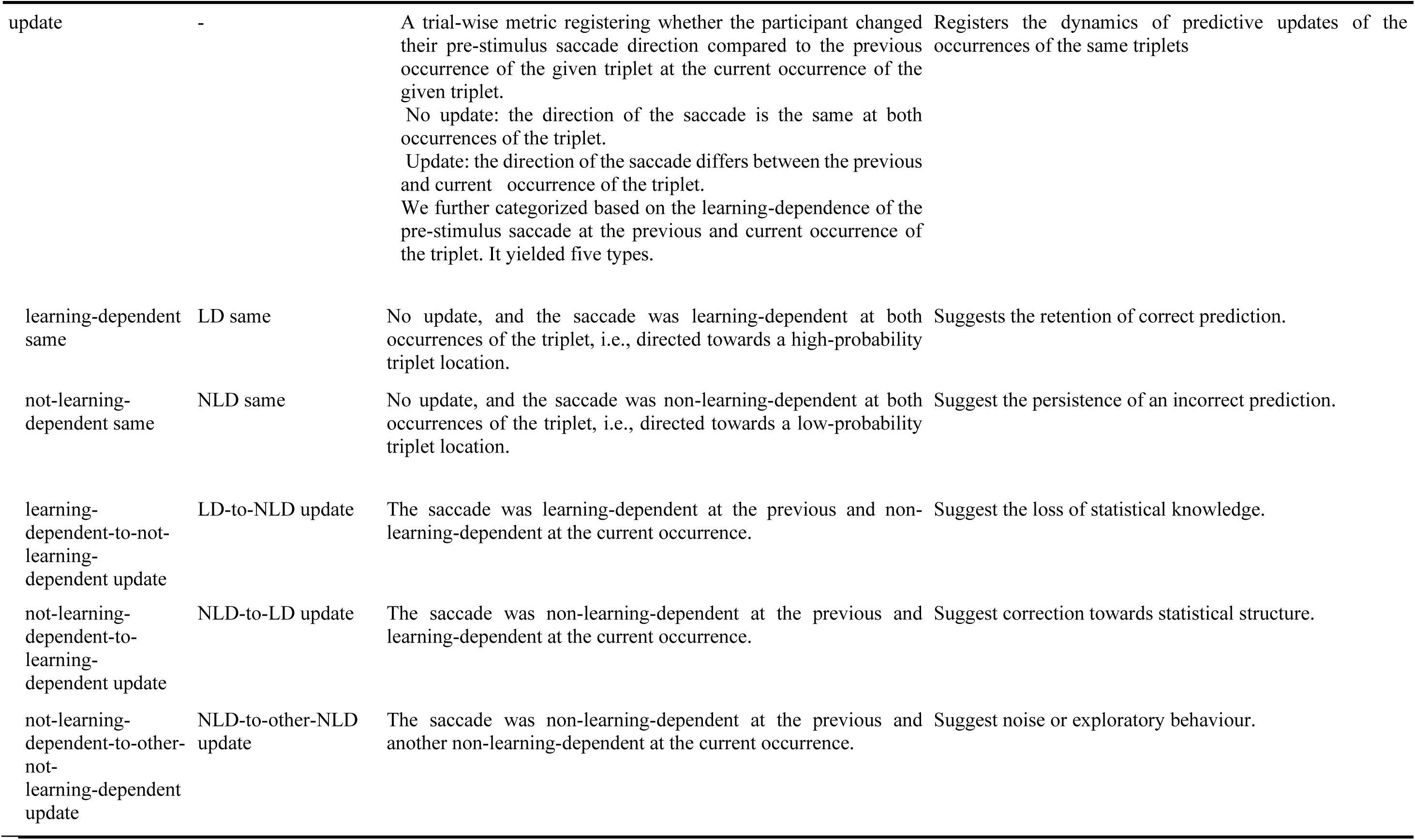
Metrics.

| Name | Abbreviation | Operationalization | Interpretation |
| --- | --- | --- | --- |
| oculomotor reaction time | oRT | The latency between the appearance of stimuli and the 100 ms fixation. | Statistical learning is defined as the decrease in oRT on the high-probability triplet compared to low-probability one. |
| saccade types | - | The first saccade during RSI, categorized based on whether the gaze of the participant was directed toward a high- or low-probability triplet location and whether this gaze was the same as the location of the upcoming stimuli. This yielded four types. | This trial-by-trial metric shows whether the anticipations are aligned with the task's statistical structure. |
| learning-dependent error | LDE | The saccade was directed toward a high-probability triplet location, but a low-probability triplet appeared. | Suggest reliance on correct knowledge but due to the probabilistic structure the prediction is incorrect. |
| learning-dependent correct | LDC | The saccade was directed toward a high-probability triplet location, and the next stimulus appeared there. | Suggest a successful prediction based on the statistical structure. |
| not-learning-dependent error | NLDE | The saccade was directed toward a low-probability triplet location, but another (low- or high-probability) triplet appeared. | Incorrect and noninformative saccade direction which does not align with the statistical structure. |
| not-learning-dependent correct | NLDC | The saccade was directed toward a low-probability triplet location and the next stimulus appeared there. | Suggest correct but not predictive behaviour since the saccade direction accidentally matched the upcoming stimuli but not the statistical structure. |
| update | - | A trial-wise metric registering whether the participant changed their pre-stimulus saccade direction compared to the previous occurrence of the given triplet at the current occurrence of the given triplet.<br>No update: the direction of the saccade is the same at both occurrences of the triplet.<br>Update: the direction of the saccade differs between the previous and current occurrence of the triplet.<br>We further categorized based on the learning-dependence of the pre-stimulus saccade at the previous and current occurrence of the triplet. It yielded five types. | Registers the dynamics of predictive updates of the occurrences of the same triplets |
| learning-dependent same | LD same | No update, and the saccade was learning-dependent at both occurrences of the triplet, i.e., directed towards a high-probability triplet location. | Suggests the retention of correct prediction. |
| not-learning-dependent same | NLD same | No update, and the saccade was non-learning-dependent at both occurrences of the triplet, i.e., directed towards a low-probability triplet location. | Suggest the persistence of an incorrect prediction. |
| learning-dependent-to-not-learning-dependent update | LD-to-NLD update | The saccade was learning-dependent at the previous and non-learning-dependent at the current occurrence. | Suggest the loss of statistical knowledge. |
| not-learning-dependent-to-learning-dependent update | NLD-to-LD update | The saccade was non-learning-dependent at the previous and learning-dependent at the current occurrence. | Suggest correction towards statistical structure. |
| not-learning-dependent-to-other-not-learning-dependent update | NLD-to-other-NLD update | The saccade was non-learning-dependent at the previous and another non-learning-dependent at the current occurrence. | Suggest noise or exploratory behaviour. |

#### Standard analysis of statistical learning

Statistical learning was defined as the difference between the reaction time of fixation on the high-probability as compared to low-probability trials. The final model contained the oculomotor reaction time as outcome variable; triplet type (high- or low-probability), epoch (1-4 as a factor), age group (children vs. adults) as fixed effect and the random intercept per participant.

#### Likelihood of saccade types

In addition to the standard ASRT measures, we implemented a novel eye-tracking-based analysis (described in Hann et al. (2026). In this approach, every trial was categorized according to the predicted stimulus (see Figure 3). First, we distinguished between learning-dependent and not-learning-dependent saccades, depending on whether the saccade was directed toward a high- or a low-probability triplet. Each trial was then classified as correct if the predicted stimulus matched the actual stimulus location, or as an error if it was directed to another stimulus. This yielded four saccade categories: learning-dependent correct (LDC), learning-dependent error (LDE), not-learning-dependent correct (NLDC), and not-learning-dependent error (NLDE). For each participant, we calculated the epoch-wise proportion of these categories, and for better interpretability, standardized them against chance-level probabilities (learning-dependent correct, LDC: 0.15625, learning-dependent error, LDE: 0.09375, not-learning-dependent correct, NLDC: 0.09375, not-learning-dependent error, NLDE: 0.65625; see Supplementary Materials and Supplementary Table S1 for details). Thus, values above one indicate performance above chance for the given trial type, while values near one reflect chance-level performance.

**Figure 3.**
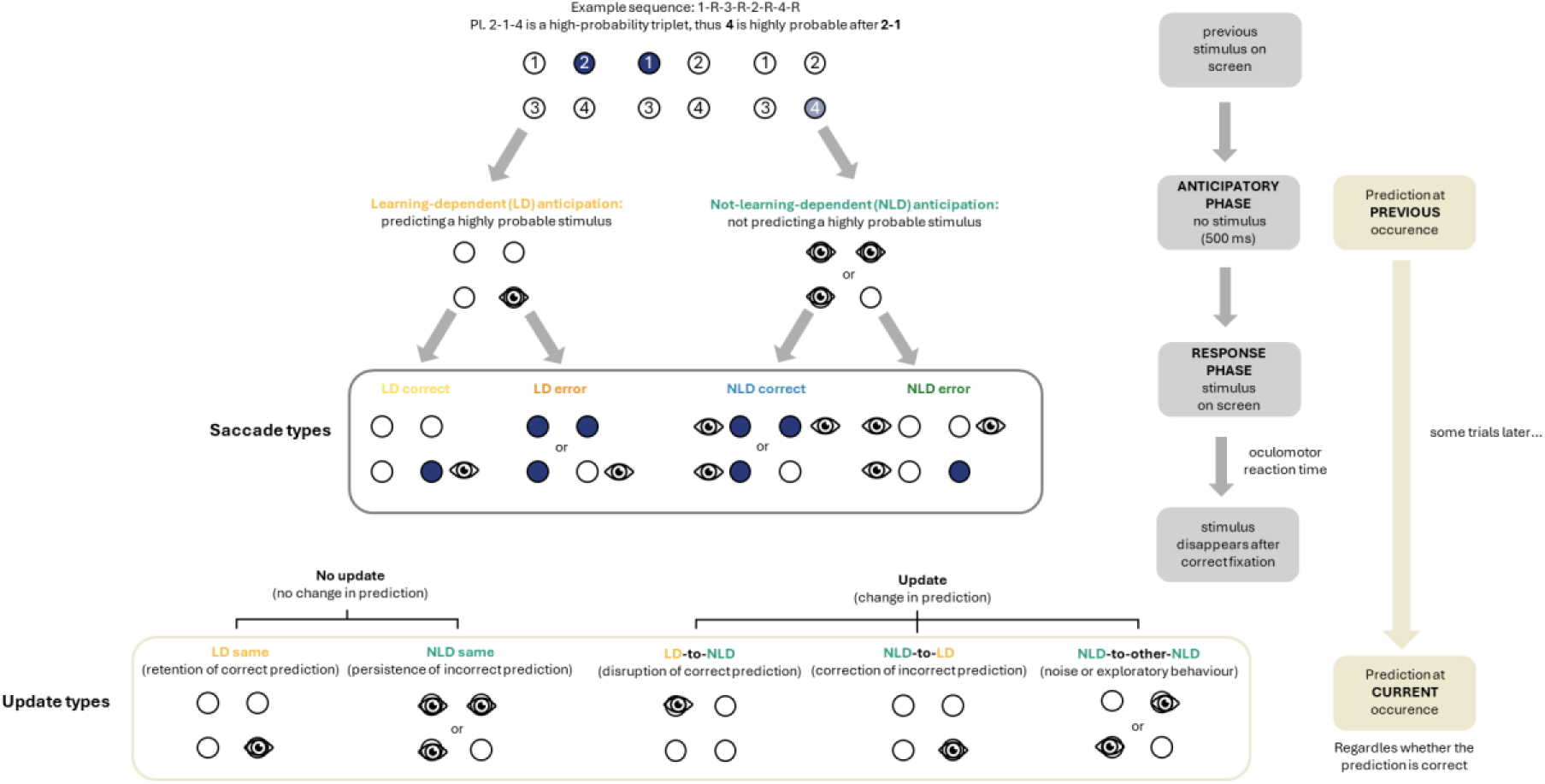
Categorization of saccade and update types. We recorded participants’ first saccade during the anticipatory phase, dividing the screen into four quadrants corresponding to the circles. Saccades toward the most likely (high-probability) stimulus location were classified as learning-dependent anticipation, others as non-learning-dependent. Trials were labelled correct if gaze matched the actual stimulus location, otherwise incorrect. This led to four categories: learning-dependent correct, learning-dependent error, non-learning-dependent correct, non-learning-dependent error. To assess updating, we examined what prediction occurred during a given triplet’s previous appearance and compared this to the prediction that occurred during the same triplet’s current appearance. Unchanged predictions were classified as no update: learning-dependent same (LD same) or non-learning-dependent same (NLD same). Changes were categorized as from learning-dependent to not-learning-dependent (LD-to-NLD) or from non-learning-dependent to learning-dependent (NLD-to-LD) or from not-learning-dependent to other not-learning-dependent (NLD-to-other-NLD).

Saccade type likelihoods were analysed by fitting a linear mixed model including epoch-wise likelihood of each saccade type as an outcome variable, epoch (1-4 as factor), saccade type (LDC, LDE, NLDC, NLDE), age group (children vs. adults) as fixed effects and the random intercept per participants.

#### Iterative updating metric

To examine group differences in prior updating, we used the iterative updating metric introduced by Hann et al. (2026). For each trial, we determined whether the given triplet had been presented before (including presentations in the first, random epoch); if not, the trial was excluded from this analysis. If the triplet had appeared previously (e.g., if the current triplet is 2-1-4, and it has occurred before), we compared the participant’s predicted stimulus in the current trial with that in its previous occurrence (e.g., the last time the participant saw the triplet 2-1-4). If the two predicted stimuli matched (e.g., the participant’s first saccade in the response-to-stimulus interval preceding the presentation of stimulus 4 in the 2-1-4 triplet was directed toward the stimulus 4 both times), we classified the trial as involving no update. If they differed (e.g., at the previous occurrence, the participant made a saccade toward the 3rd circle but now they made one toward the 4th one), the trial was classified as an update. This procedure resulted in a binary variable, coded as 0 for no update and 1 for update.

Using this measurement, we ran a generalized linear mixed model with the update likelihood as an outcome variable, fixed effects included epoch (1-4 as a factor), age group (children vs. adults) and the random intercept per participant.

#### Update type likelihoods

The iterative update metric indicates only whether a participant changed their prediction compared to the previous occurrence of the same triplet. To gain deeper insight, we further analysed how responses changed from one occurrence to the next. A shift from a learning-dependent to a not-learning-dependent response (LD-to-NLD update) may reflect disruption of previously acquired knowledge. Conversely, a shift from not-learning-dependent to learning-dependent (NLD-to-LD update) suggests correction of the participant’s internal model of environmental regularities. Updates from one not-learning-dependent direction to another (NLD-to-other-NLD) may reflect noise or exploratory behaviour. In trials without an update, participants could either persist with a not-learning-dependent response (NLD same), indicating no evidence of new learning, or maintain a learning-dependent response (LD same), reflecting stability in their acquired knowledge. Similarly to the trial type likelihoods, first we calculated the proportion of these update types in each epoch for each participant. Then, update type likelihoods were calculated by these proportions against their chance-level probabilities (LD-to-NLD: 0.1875, NLD-to-LD: 0.1875, NLD-to-other-NLD: 0.375, NLD same: 0.1875, LD same: 0.0625, for a detailed description of these probabilities, see Supplementary Materials and Supplementary Table S2).

These categories were entered into a linear mixed model with the update type likelihood as an outcome variable. Fixed effects included update type (LD-to-NLD / NLD-to-LD / NLD-to-other-NLD / NLD same / LD same), epoch (1-4 as a factor), and age group (children vs. adults), with a random intercept and random slope for epoch per participant (with correlation).

## Results

### Do children and adults differ in the classic ASRT metrics?

#### Oculomotor reaction times

To examine developmental differences in statistical learning using the standard analysis, we fitted a linear mixed-effects model predicting oRT from Triplet type (high vs. low probability) × Epoch (1–4) × Group (children vs. adults), with a random intercept for subjects, see Figure 4. There was a significant main effect of Triplet type, *F*(1, 81480.71) = 242.45, *p* < 0.001, indicating overall statistical learning, with faster oculomotor responses to high-than low-probability trials (*high–low = −30.30 ms, SE = 1.94 ms, p < 0.001*). A significant main effect of Epoch, *F*(3, 81480.35) = 43.63, *p* < 0.001, showed a gradual, general slowing across the task (*epoch 1 - epoch 2: −15.17 ms, SE = 2.75 ms, p < 0.001; epoch 2 - epoch 3: −6.39 ms, SE = 2.75 ms, p = 0.093; epoch 3 - epoch 4: −8.94 ms, SE = 2.75 ms, p = 0.006*). The main effect of Group was also significant, *F*(1, 66.13) = 10.76, *p* = 0.002, reflecting overall differences in general oculomotor response speed between children and adults, with children being slower with an overall 50.60 ms (*SE = 15.40 ms, p = 0.001*). The interaction between triplet type and epoch did not reach significance, *F*(3, 81481.02) = 1.86, *p* = 0.134, indicating no significant evidence that statistical learning changed across epochs.

**Figure 4.**
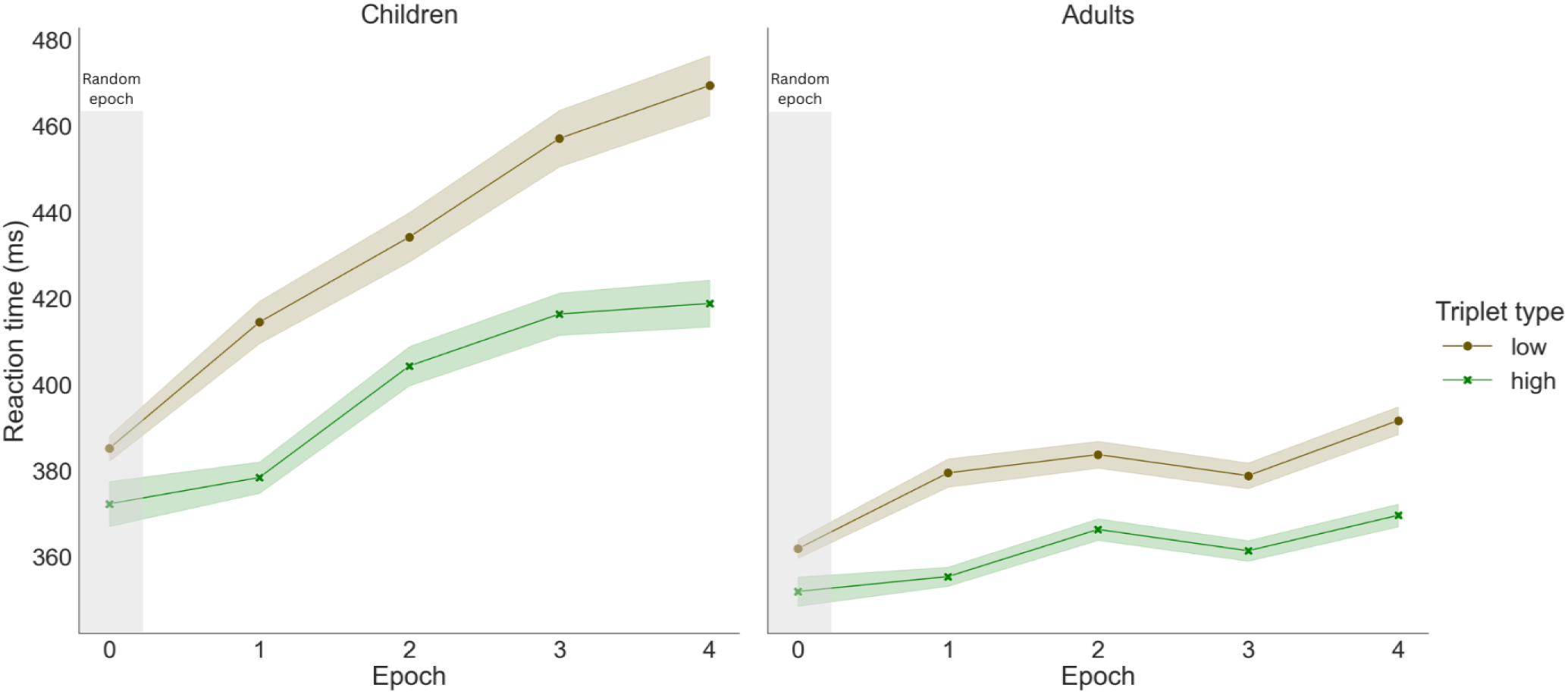
Oculomotor reaction time in children (left figure) and adults (right figure), by epochs. The brown color with circle markers show the oculomotor RT of low-probability triplets, the green color with x markers the oculomotor RT of high-probability triplets. The gap between the two lines indicates the magnitude of statistical learning. We found a significantly greater amount of statistical learning in children than in adults.

The Triplet type × Group interaction was significant, *F*(1, 81480.71) = 24.57, *p* < 0.001. Post hoc tests revealed that both groups showed learning as indicated by the high–low probability triplet differences (*children: −39.90 ms, SE = 2.98 ms, p < 0.001; adults: −20.60 ms, SE = 2.50 ms, p < 0.001*), with children showing a greater amount of statistical learning (*high–low difference in children vs. adults: −19.30 ms, SE = 3.89 ms, p < 0.001*). The Epoch × Group interaction was also significant, *F*(3, 81480.35) = 20.52, *p* < 0.001, suggesting that the groups differed in general oculomotor speed only from the second epoch on (*epoch 1, children - adults: 28.90 ms, SE = 15.80 ms, p = 0.068; epoch 2, children - adults: 43.90 ms, SE = 15.80 ms, p = 0.005; epoch 3, children - adults: 66.70 ms, SE = 15.80 ms, p < 0.001; epoch 4, children - adults: 63.00 ms, SE = 15.80 ms, p < 0.001*). Finally, the three-way interaction of Triplet type × Epoch × Group was not significant, *F*(3, 81481.02) = 1.30, *p* = 0.272, suggesting that the groups did not significantly differ in how statistical learning evolved over time.

### Are children more likely to commit learning-dependent errors than adults?

To examine whether children and adults differed in the extent to which their errors reflected acquired statistical knowledge, we analysed the likelihood of different saccade types across development. A higher likelihood for the learning-dependent saccade types reflects a stronger statistical learning effect. We fit a linear mixed model with a random intercept for subjects, examining the likelihood of saccade Types (LDC, LDE, NLDC, NLDE) across Epochs (1-4) and Groups (children vs. adults), see Figure 5. There was a significant main effect of Type, *F*(3, 990) = 122.41, *p* < 0.001, indicating differences in likelihood across saccade types. Post hoc pairwise comparisons revealed that learning-dependent errors were more likely than any other saccade types (difference from: *learning-dependent correct: 0.13, SE = 0.02, p < 0.001, not-learning-dependent correct: 0.28, SE = 0.02, p < 0.001, not-learning-dependent error: 0.28, SE = 0.02, p < 0.001*). The second most likely were the learning-dependent correct saccades, being significantly more likely than the not-learning-dependent correct (*0.16, SE = 0.02, p < 0.001*), and error (*0.15, SE = 0.02, p < 0.001*) saccades. There was no difference between not-learning-dependent correct and not-learning-dependent error saccade types (*0.01, SE = 0.02, p = 0.986*).

**Figure 5.**
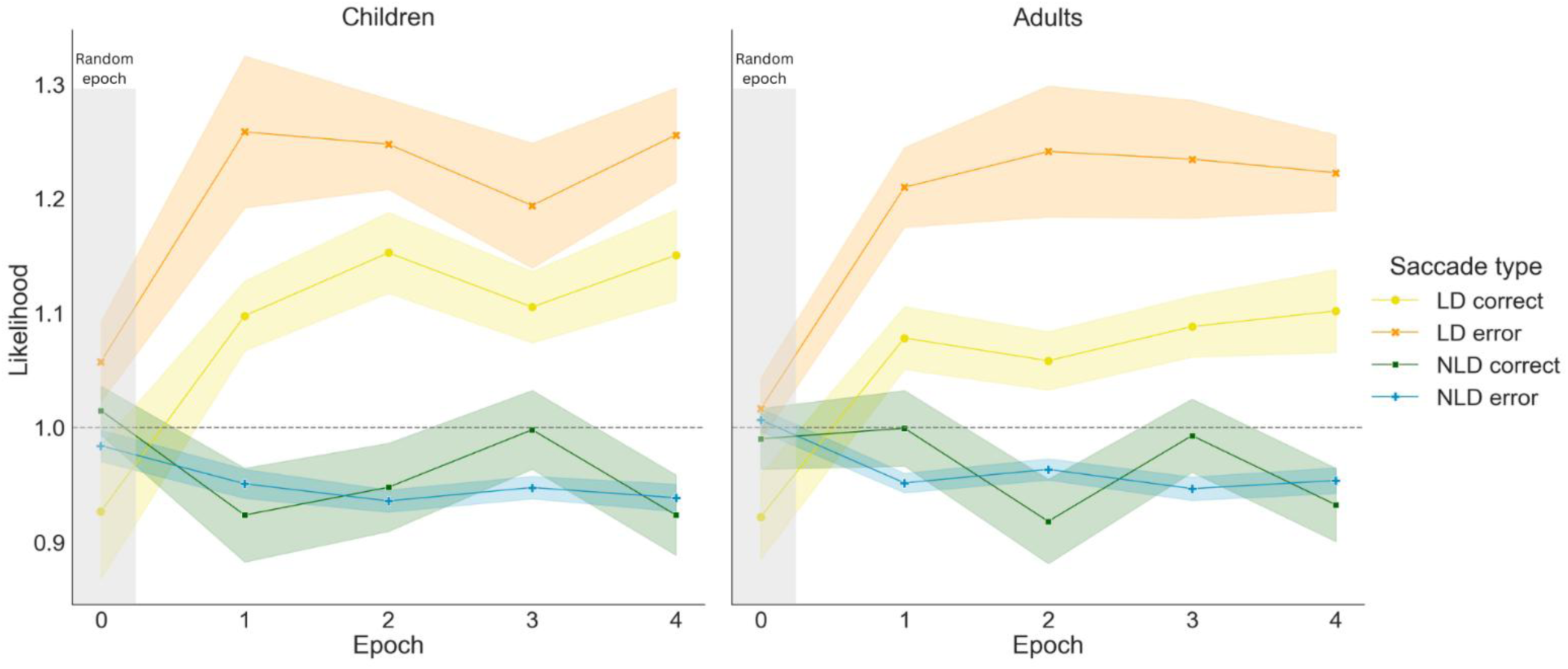
Saccade type likelihoods in children (left) and adults (right). The y-axis represents the likelihood, the x-axis shows time measured in epochs, and the lines represent the learning-dependent correct (LDC, yellow), learning-dependent error (LDE, orange), not-learning-dependent correct (NLDC, green), not-learning-dependent error (NLDE, blue) saccade types. The error bands represent standard error of the mean.

No other effects were significant. Either the main effect of Epoch, *F*(3,990) = 0.04, *p* = 0.990, or the main effect of Group, *F*(1,66) = 0.46, *p* = 0.501, was not significant, indicating no significant changes in overall likelihood across epochs or between groups. The Type × Epoch interaction also did not reach significance, *F*(9, 990) = 0.77, *p* = 0.646, indicating that the likelihood of different saccade types did not show significantly different trajectory throughout the task. The non-significant Type × Group interaction, *F*(3, 990) = 1.18, *p* = 0.316, shows a similar likelihood of different saccade types in the age groups, while the non-significant Epoch × Group interaction, *F*(3, 990) = 0.34, *p* = 0.798, suggests that the likelihood changes did not significantly depend on the age group. The three-way interaction between Type × Epoch × Group was also not significant, *F*(9, 990) = 0.58, *p* = 0.812, as children and adults did not significantly differ in how likelihoods of the saccade types changed as the task progressed.

### Do children update their responses more than adults?

To investigate developmental differences in whether participants tend to stick to their previous predictions or rather update it, a generalized linear mixed model was conducted to examine the effects of Epoch (1-4) and Group (children vs. adults) on update ratio compared to trials with no update, see Figure 6. There was a significant main effect of Group, χ²(1) = 10.21, *p* = 0.001, indicating overall differences between children and adults in the updating ratio. Post hoc pairwise comparisons revealed higher update ratio in children (log-odds 0.285, SE = 0.084, *p* < 0.001). The main effect of Epoch was not significant, χ²(3) = 1.26, *p* = 0.739, meaning that the update ratio was stable across epochs. The Epoch × Group interaction was also not significant, χ²(3) = 1.77, *p* = 0.622, meaning that the changes in the update ratio did not significantly differ between the two age groups as the task progressed.

**Figure 6.**
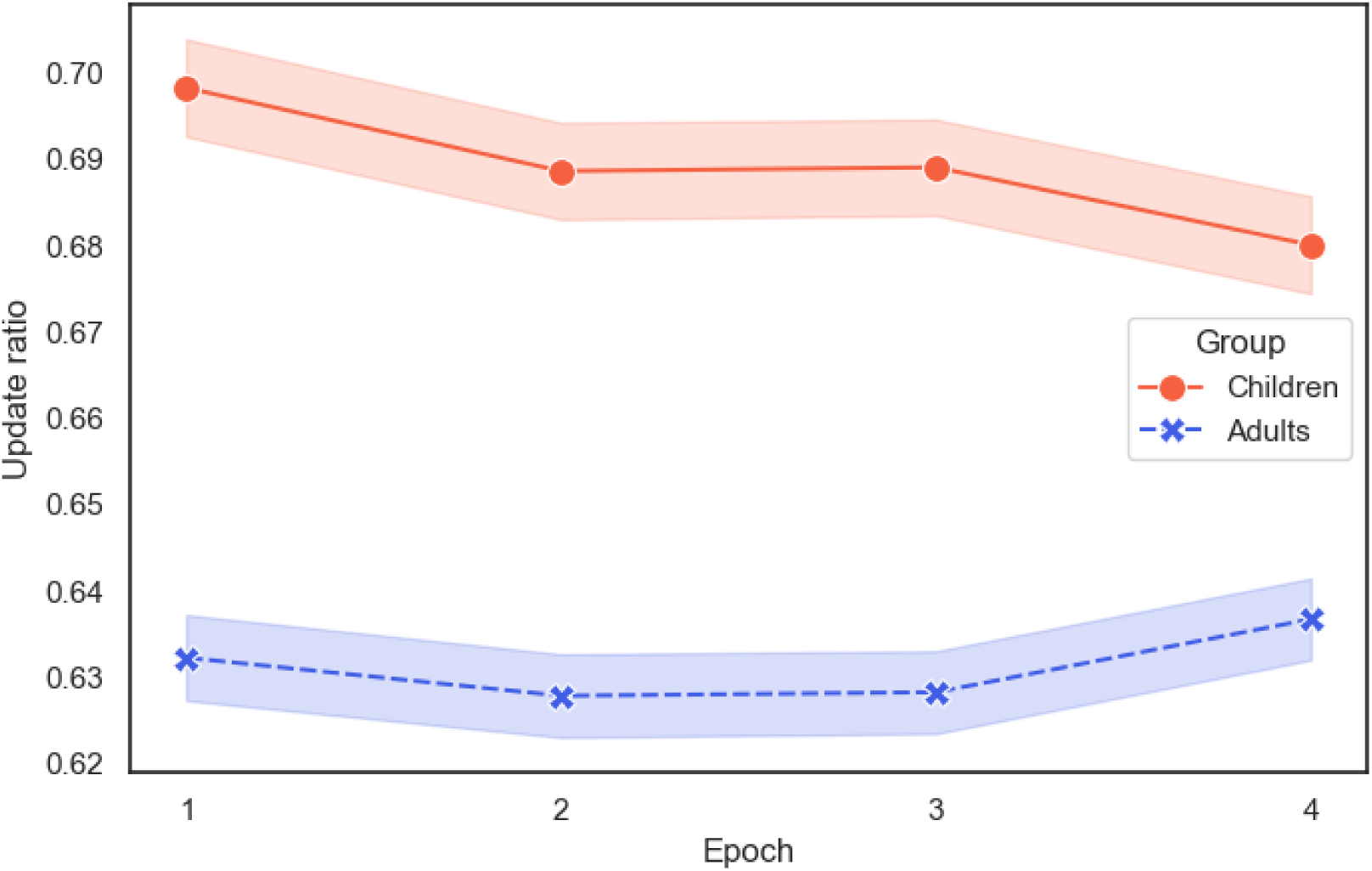
The ratio of updates in the age groups. The y-axis shows the ratio of updates compared to all trials, the x-axis represents time as measured in epochs. The lines represent the age groups: children (red, with circular marker) and adults (blue, with x marker). Error bands represent standard errors of the mean.

### Is there a developmental difference in updating?

To further understand the mechanisms underlying developmental differences in belief updating, we examined whether the tendency to update responses depended on the type of prediction made on the previous trial. A generalized linear mixed model was conducted to test the effects of Epoch (1-4), Previous saccade type (learning-dependent correct, not-learning-dependent correct, learning-dependent error, not-learning-dependent error), and Group (children vs. adults) on Update (0/1). There was a significant main effect of Previous saccade type, χ²(3) = 24.24, *p* < 0.001. Post hoc pairwise comparisons showed that learning-dependent correct did not significantly differ from learning-dependent error (*p* = 0.503), and not-learning-dependent correct did not differ from not-learning-dependent error (*p* = 0.947). However, both learning-dependent types (learning-dependent correct and learning-dependent error) differed significantly from both not-learning-dependent types (all *ps* < 0.001), with higher update likelihood following not-learning-dependent saccades compared to learning-dependent responses. This together means that participants updated their responses less frequently when the previous response aligned with the underlying structure of the task vs. when it did not align, regardless of whether it was an error or not.

The main effects of Epoch, χ²(3) = 1.06, *p* = 0.788, and Group, χ²(1) = 2.31, *p* = 0.129, were not significant. The interactions between Epoch and Previous response type, χ²(9) = 10.18, *p* = 0.336, Epoch and Group, χ²(3) = 1.19, *p* = 0.755, Previous response type and Group, χ²(3) = 1.68, *p* = 0.642, and the three-way interaction, χ²(9) = 7.26, *p* = 0.610, were also not significant.

### How do children and adults update their responses?

To further characterize developmental differences in response updating, we examined the specific types of transitions participants made between predictions of consecutive occasions of the same triplet. A linear mixed model was conducted to examine the effects of Epoch (1-4), Update type (LD same, NLD same, LD-to-NLD, NLD-to-LD, NLD-to-other-NLD), and Group (children vs. adults) on the likelihood of update types, see Figure 7. There was a significant main effect of Update type, *F*(4, 1254) = 324.57, *p* < 0.001. Post hoc comparisons showed that all update types differed significantly from each other (*p*s < 0.001), except for NLD-to-LD and LD-to-NLD (*p* = 0.972). Overall, LD same had the highest likelihood, while NLD-to-other-NLD had the lowest. Taken together, this suggests that participants were most likely to repeat predictions that aligned with the task probabilities. However, they also showed a general tendency to repeat rather than switch their predictions even when those predictions did not align with the task structure. Notably, switches from a not-learning-dependent to a learning-dependent prediction were no more likely than switches in the opposite direction.

**Figure 7.**
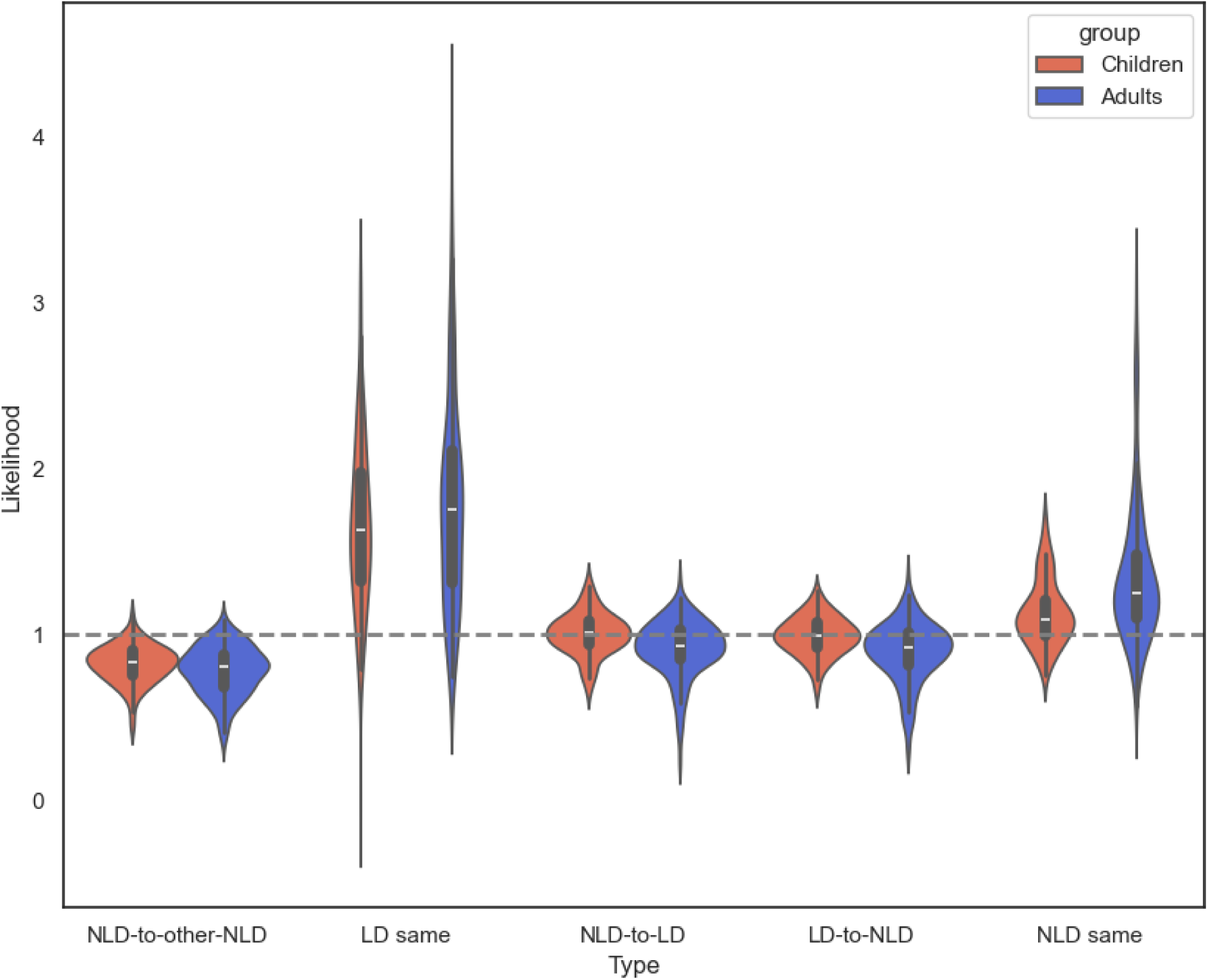
Likelihood of update types in children (red color, on the left) and adults (blue color, on the right). The y-axis represents the likelihood of update types, the x-axis shows the update types. The dashed grey line represents chance level.

There was a significant Update type × Group interaction, *F*(4, 1254) = 12.77, *p* < 0.001. Post hoc comparisons examined group differences within each update type. There was no significant group difference for NLD-to-other-NLD (0.04, SE = 0.04, *p* = 0.310). However, children showed lower likelihood than adults for LD same (−0.136, SE = 0.04, *p* = 0.001) and NLD same update types (−0.224, SE = 0.04, *p* < 0.001), and higher likelihood than adults for NLD-to-LD (0.089, SE = 0.04, *p* = 0.029) and LD-to-NLD (0.097, SE = 0.04, *p* = 0.018) update types. This suggests that children were less likely to repeat their previous prediction than adults and were more likely to switch to a new prediction.

## Discussion

Our study provides new evidence that children extract environmental patterns more effectively than adults, even when measured with sensitive eye-tracking methods. Consistent with earlier reports on manual reaction times, children showed stronger statistical learning in their oculomotor reaction times, but the analysis of saccadic eye-movements revealed an even richer picture. Children showed a higher tendency to update their responses than adults, suggesting greater behavioural flexibility. At the same time, children were less likely to repeat the same saccade type and more likely to switch between learning-dependent and non-learning-dependent predictions compared to adults. This pattern indicates that children adopt a less stable predictive strategy, whereas adults show more consistency in maintaining learned predictions – with the former strategy resulting in better statistical learning. Strikingly, the fundamental capacity to process and apply prediction errors remained remarkably invariant across development.

One of the candidate mechanisms that might underlie the developmental differences in statistical learning is the computation and weighting of prediction errors. Evidence from the reinforcement learning literature indicates that developmental differences are less pronounced in error computation itself, and instead emerge in how these errors are used to guide behaviour (Van Den Bos et al., 2012; Zhang et al., 2019). However, statistical learning paradigms such as the ASRT differ from reinforcement learning in that learning is driven by exposure to probabilistic regularities rather than explicit rewards. This distinction may explain why we did not observe any error-related developmental differences. Prior findings have linked age-related differences to the maturation of reward-sensitive neural systems, including the striatum and prefrontal cortex, suggesting that the age-related difference lies more in reward processing than in predictive processing per se (Van Den Bos et al., 2012). Our results support this view, suggesting similarly calculated and processed errors in children and adults. Alternatively, it is possible that statistical learning is not error-driven at all, thus, the lack of age difference on the errors just reflect that it is an irrelevant factor in this type of learning (Hann et al., 2026). Future studies should address this question directly, by comparing children and adults on various tasks with and without rewards.

In our study, we found that children tend to update their responses more than adults. Specifically, they are more likely to switch from learning-dependent to not-learning-dependent anticipations and from not-learning-dependent to learning-dependent anticipations than adults. The former suggests the loss of statistical knowledge (as they update from a prediction that aligns with the task regularity to a prediction that does not), while the latter suggests correction towards the statistical structure (as they update from a non-task-aligned prediction to a task-aligned one). In contrast, adults tend to update less and stick to their previous choices, even when this means persistence of an incorrect prediction (i.e., no update on a not-learning-dependent triplet). More switching between predictions may reflect developmental differences in information processing style, with children showing a broader way to sample the world (Frank et al., 2021; Wan & Sloutsky, 2025). This may relate to reduced working memory capacity in childhood that may limit the stable maintenance of previously accumulated evidence, effectively promoting repeated sampling of the environment (Wan & Sloutsky, 2025). However, simply forgetting previous predictions could suggest that children do not retain the regularity itself either, which was not the case in our study, as children showed better statistical learning.

From a Bayesian perspective, the combination of similar error processing and age-related differences in belief updating is highly informative. As the updating of the prior belief depends on the combination of the weighted priors and the weighted sensory stimulus (i.e., how much trust the learner puts into each of these) (Moran et al., 2013), similar error signals coupled with differential updating imply a shift in the balance of the prior and sensory stimulus weight. Specifically, greater updating in children may suggest a relative down-weighting of prior precision (or up-weighting of sensory precision), resulting in a stronger influence of incoming information on belief updating. Furthermore, relating these findings to the reinforcement learning literature offers an additional perspective. The observed pattern is consistent with accounts of choice-confirmation bias documented in reinforcement learning, whereby previously selected options are preferentially reinforced, independent of their objective value (Talluri et al., 2018). Within this framework, adults may overweight evidence that is consistent with prior choices while underweighting disconfirming information, leading to more stable but less flexible behaviour. Our results extend this pattern to a non-reward context: even in the absence of reward, adults appear more likely to persist with prior choices. Taken together with the absence of developmental differences in error processing, it appears that the developmental trajectory of non-rewarded statistical learning resembles reinforcement learning. While both domains may share similar updating dynamics and confirmation biases, they appear to diverge in the role of error signals. However, future studies should directly compare these two forms of learning within a unified paradigm, particularly given that similar underlying mechanisms may give rise to divergent behavioural outcomes (e.g., adults outperforming children in reinforcement learning, but not in statistical learning).

The trajectory of statistical learning shows a varied picture across modalities, types of probability, and measurement methods, which might suggest that children’s strategies may not be equally well suited to all statistical learning tasks. In light of the observed higher oculomotor learning in children, it might be concluded that in the ASRT task, higher updating rates (which may be considered as exploration) can be beneficial (Hann et al., 2026; Pesthy et al., 2023). The task involved a non-adjacent sequence embedded in noise, comprising 64 triplets (16 high-probability and 48 low-probability). Within this context, children’s strategy to explore the task might result in a broader – although potentially less stable – knowledge, whereas adults stabilize their predictions at an earlier stage. In contrast, for linguistic stimuli, learning reaches adult levels as early as childhood and does not develop further, which might be explained by an innate tendency to pay attention to these stimuli (Forest et al., 2023). This might also be the case in tasks where the stimuli to be learned are signalled, such as the contextual cueing task (Merrill et al., 2013). When acquiring deterministic or adjacent probabilities, that is, when the next stimulus can be predicted with a probability of one by the immediately preceding stimulus, exploration is not beneficial as the regularities are easily obtained. Rather, exploration might result in losing the information that needs to be learned. Accordingly, children’s exploratory learning strategy may confer no advantage (developmental invariance) or even a disadvantage (age-related improvement) in these tasks (Arciuli & Simpson, 2011; Bertels et al., 2014; Jost et al., 2015; Lukács & Kemény, 2015; Lum et al., 2010; Raviv & Arnon, 2018; Schlichting et al., 2017; Shufaniya & Arnon, 2018). In tasks using offline measurement, other cognitive processes, such as metacognitive abilities may also influence the results (Siegelman, Bogaerts, Christiansen, et al., 2017). As these processes are still developing in childhood, they may further contribute to apparent age-related disadvantages.

Various frameworks have proposed that internal models can constrain statistical learning during development (Forest et al., 2023; Janacsek et al., 2012; Seitz, 2021; R. Wu et al., 2016). The competition model proposed by Janacsek et al. (2012) suggested that in childhood, we detect raw probabilities in the environment more easily as our internal models are not yet fully developed. As we age, our internal models develop and function as a competition to newly acquired regularities. Thus, previously acquired knowledge might bias the learning of new non-adjacent probabilistic information, making statistical learning on the ASRT task less effective in adolescence and adulthood than in childhood (Janacsek et al., 2012). Our results align with these accounts: we found better statistical learning in children than adults as well as signs of more exploration in childhood when investigating the update types. Specifically, children were more likely to switch from learning-dependent to not-learning-dependent triplets and vice versa, while adults were more likely to repeat their previous responses, indicating a stronger tendency to exploit established predictions. This pattern is consistent with the developmental shift in explore/exploit trade-off (Frankenhuis & Gopnik, 2023; Gopnik, 2020). In detail, this framework suggests that humans regulate a trade-off between exploring unknown aspects of the environment and exploiting known rewards, with a bias toward exploration in childhood that shifts toward more exploitative, goal-directed behaviour in adulthood. Similarly, Wu et al. (2016) differentiate between input-driven and knowledge-driven learning, arguing that development depends more on input than prior knowledge. In childhood, limited experience reduces the influence of existing knowledge, allowing learning to be guided mainly by environmental patterns. In adulthood, however, learning is increasingly shaped by prior knowledge, such as schemas, internal models, routines, and assumptions (Frank et al., 2021; Seitz, 2021; R. Wu et al., 2016). It is important to note that our study was not designed to test these frameworks. Nevertheless, our results are in line with the predictions of these frameworks and highlight the importance of examining the developmental trajectory of statistical learning in greater detail.

Besides age, several other factors might influence statistical learning development and exploration. One key factor might be socio-economic status, more specifically, childhood unpredictability (Farkas et al., 2026; Xu et al., 2023). Xu et al. (2023) showed that children who experienced their environments as unpredictable explored less and were more likely to repeat their previous choices in a reinforcement learning task. In a study employing the motor ASRT task in adults, subjective current socio-economic status had negative effect on statistical learning, while childhood unpredictability was associated with better statistical learning, as reflected in the faster attainment of asymptotic performance during the learning phase (Farkas et al., 2026). At first glance, these studies seem contradictory: childhood unpredictability is associated with less exploration but better statistical learning and as we discussed above, statistical learning might benefit from higher exploration which might explain children’s advantage on the task. However, there are two important factors to consider: (1) the presence of reward and (2) the type of exploration in reinforcement and statistical learning tasks. In reinforcement learning tasks, exploration is needed in a goal-directed, reward-based way. In contrast, statistical learning is unsupervised and relies more on passive, input-driven extraction of regularities. Thus, children growing up in unpredictable environments may engage less in strategic, reward-based exploration, while still showing heightened sensitivity to environmental patterns, leading to enhanced statistical learning.

In sum, our findings demonstrate that children’s advantage in statistical learning is not driven by differences in error processing, but is instead closely linked to how information is used to update behaviour. Across analyses, children showed a more flexible, exploratory response pattern, whereas adults exhibited greater stability and persistence in their choices, even at the cost of reduced sensitivity to probabilistic structure. These results suggest that developmental differences in statistical learning reflect a shift in the balance between exploration and exploitation, potentially grounded in differences in the weighting of prior knowledge and incoming sensory information. By combining oculomotor measures with computationally informed interpretations, the present study provides a more fine-grained account of how learning unfolds across development. Future work should directly compare statistical and reinforcement learning within unified paradigms and further investigate how task characteristics, environmental factors, and internal models interact to shape learning strategies across the lifespan.

## Data and code availability

The behavioral datasets generated and analyzed during the current study, as well as the custom analysis code, are available in the OSF repository at https://osf.io/qm8av/.

## Acknowledgement

This work was supported by the French National Research Agency (ANR-24-CE37-5807), the National Brain Research Program (NAP2022-I-2/2022), and the Hungarian Scientific Research Fund (NKFIH ADVANCED153150), all awarded to D.N.

## Declaration on the Use of AI-Assisted Technologies

The authors, who are not native English speakers, used an AI-assisted tool to improve the readability of author-written text. The tool was not used to generate scientific content or ideas. All concepts, analyses, and conclusions are the authors’ own, and the authors take full responsibility for the final manuscript.

