## Supplementary Material for "Flexible belief updating drives the childhood advantage in statistical learning"

### Supplementary Methods

#### Baseline probabilities of saccade types

We calculated the chance-level probabilities of each saccade type based on the chance-level response probability for each stimulus location (P = 0.25 for each of the four circles), and the chance-level probability of all stimulus outcomes (high-probability triplets: P = 0.625, and each of the three low-probability triplet options: P = 0.125). The calculated probabilities are shown in Supplementary Table S1.

**Supplementary Table S1.** Baseline (chance-level) probabilities of each saccade type.

| Saccade direction | Actual outcome | Correct | P of event | Label |
|---|---|---|---|---|
| High-prob (P = 0.25) | High-prob (P = 0.625) | Yes | 0.15625 | LDC |
| | Low-prob/1 (P = 0.125) | No | 0.03125 | LDE |
| | Low-prob/2 (P = 0.125) | No | 0.03125 | LDE |
| | Low-prob/3 (P = 0.125) | No | 0.03125 | LDE |
| Low-prob/1 (P = 0.25) | High-prob (P = 0.625) | No | 0.15625 | NLDE |
| | Low-prob/1 (P = 0.125) | Yes | 0.03125 | NLDC |
| | Low-prob/2 (P = 0.125) | No | 0.03125 | NLDE |
| | Low-prob/3 (P = 0.125) | No | 0.03125 | NLDE |
| Low-prob/2 (P = 0.25) | High-prob (P = 0.625) | No | 0.15625 | NLDE |
| | Low-prob/1 (P = 0.125) | No | 0.03125 | NLDE |
| | Low-prob/2 (P = 0.125) | Yes | 0.03125 | NLDC |
| | Low-prob/3 (P = 0.125) | No | 0.03125 | NLDE |
| Low-prob/3 (P = 0.25) | High-prob (P = 0.625) | No | 0.15625 | NLDE |
| | Low-prob/1 (P = 0.125) | No | 0.03125 | NLDE |

| | | | | |
|---|---|---|---|---|
| | Low-prob/2 (P = 0.125) | No | 0.03125 | NLDE |
| | Low-prob/3 (P = 0.125) | Yes | 0.03125 | NLDC |
| Total LDC | | | 0.15625 | |
| Total LDE | | | 0.09375 | |
| Total NLDC | | | 0.09375 | |
| Total NLDE | | | 0.65625 | |
| Total | | | 1 | |

*Note*. High-prob = the last element of a high-probability triplet, low-prob/1, 2, or 3: the last element of one of the possible low-probability triplets. Ps reflect the chance-level probability of a given outcome. LDC = learning-dependent correct, LDE = learning-dependent error, NLDC = not-learning-dependent correct, NLDE = not-learning-dependent error.

**Baseline probabilities of update types**

We calculated the update type chance-level probabilities as follows. We computed the joint probability of (a) a given saccade type to happen (see above in Table S1), (b) the current triplet to occur again, and (c) a specific saccade type occurring at the current trial. This resulted in the chance-level probabilities demonstrated in Table S2.

**Supplementary Table S2.** Baseline (chance-level) probabilities of each update type.

| **Update type** | P |
|---|---|
| LD same | 0.0625 |
| NLD same | 0.1875 |
| LD-to-NLD update | 0.1875 |
| NLD-to-LD update | 0.1875 |
| NLD-to-other-NLD update | 0.375 |

*Note.* P refers to the baseline (chance-level) probability of each event.

### Supplementary Results

**Supplementary Table S3.** Model details for all models

| Model | ICC | Marginal $R^2$ / Conditional $R^2$ |
|---|---|---|
| **Oculomotor reaction time** | 0.05 | 0.012 / 0.064 |
| **Saccade type likelihood** | 0 | 0.263 / 0.264 |
| **Update** | 0.04 | 0.006 / 0.043 |
| **Update types** | 0 | 0.519 / 0.521 |

**Supplementary Table S4.** Pairwise contrasts of the epoch main effect in the RT ~ triplet type x epoch x group model, with a random intercept for subject

| Contrast | Estimate | SE | df | z-ratio | p-value |
|---|---|---|---|---|---|
| epoch1 - epoch2 | -15.17 | 2.75 | Inf | -5.514 | <.0001 |
| epoch1 - epoch3 | -21.55 | 2.75 | Inf | -7.846 | <.0001 |
| epoch1 - epoch4 | -30.49 | 2.75 | Inf | -11.086 | <.0001 |
| epoch2 - epoch3 | -6.39 | 2.75 | Inf | -2.325 | 0.0925 |
| epoch2 - epoch4 | -15.32 | 2.75 | Inf | -5.57 | <.0001 |
| epoch3 - epoch4 | -8.94 | 2.75 | Inf | -3.252 | 0.0063 |
